# Third Backcross Generation Indonesian Indigenous Chicken *Kampong* Broiler-Type (*Kambro*) *SOX5* Gene Polymorphism

**DOI:** 10.1101/2021.02.17.431737

**Authors:** T Ardo, A. B. I. Perdamaian, I. W. S. Mahardhika, B S Daryono

## Abstract

The comb is an accessory organ on the head of chicken which is influenced by testosterone hormone and can be used as an indicator of chicken’s fertility. Comb shape is related to climate adaptation and associated with a dominant mutation in chicken chromosome 1. Therefore this research was aimed to study the association between *Pea-comb* shape and *SOX5* gene polymorphism in the population of progenies (BC-III *Kambro*) derived from a crossbreed between females *Pelung* and males second backcross generation *Kampong* Broiler-Type (BC-II *Kambro*). Chicken *(Gallus gallus) SOX5* gene was acquired from NCBI GenBank with the Ref. Seq. **418195**. Primers used to amplify the *SOX5* gene are `(F):5`’-AGGTAGCCATGGTGACAAGC-3’`,(R):5`’`-`GATCTGTGAGGCAGCCAGTT-3’. Progenies showed 100% *Pea-comb* shape, while parental generation composed of *Pea-comb* shape and *Single-comb* shape. PCR-RFLP and endonuclease restriction enzyme *HindIII* were unable to determine the genotype of the female parent with *Pea-comb*. The result of *SOX5* gene polymorphism showed the comb shape uniformity between progenies and parental of *Kambro* backcross generation. This study concluded that the genotype of *Pea-comb* shape female *Pelung* was undetermined and there was no polymorphism of the *SOX5* gene between pea and single comb. PCR-RFLP using endonuclease restriction enzyme *HindIII* produced both target products and non-target/artifact products. The sequencing procedure was required to provide nucleotide sequences.

## 1. Introduction

In 2012, Indonesian indigenous chicken meat production only covers 16% of the total Indonesia chicken meat production, while imported broiler-type chicken meat provided the rest. This cause by the disproportionate production between indigenous chicken and imported broiler-type. To tackle this issue, the Faculty of Biology Universitas Gadjah Mada step-up to innovate by the selective breeding program of *Pelung* chicken.

In this study, we exploited the genetic resources of *Pelung* chicken from Cianjur, West Java, Indonesia. As one of 34 Indonesian indigenous chicken, *Pelung* chicken has a higher body weight growth, unique meat flavor and superior posture compare with other indigenous breeds [1]. Although *Pelung* has diverse characteristics, high variation in body weight, slow growth performance, and lower reproductive traits are obstacles for its commercialization. *Kampong* Broiler-Type (*Kambro*) chicken line selective breeding program was conducted to increase the performance of *Pelung* by crossing with Broiler Cobb 500 [1].

Indonesian indigenous chicken breed known as *Ayam Kampong* has high nutritional and has high demand in the local market. *Kampong* chicken is mostly bred by villagers in Indonesia because it is easily maintained, has high nutritional meats, strong posture, and *Ayam Kampong*’s eggs are also in high demand due to its nutritional content. Indonesian indigenous chicken into 34 distinct breeds, *Ayunai, Balenggek, Banten, Bangkok, Burgo, Bekisar, Cangehgar, Cemani, Ciparage, Gaok, Jepun, Kampung, Kasintu, Kedu (hitam* and *putih), Pelung, Lamba, Maleo, Melayu, Merawang, Nagrak, Nunukan, Nusa Penida, Olagan, Rintit* or *Walik, Sedayu, Sentul, Siem, Sumatera, Tolaki, Tukung, Wareng, Sabu*, and *Semau* [2].

A trend of exploiting indigenous chicken breeds into the local poultry sector has been increasing especially in developing countries, for example, a study on Korean native chicken [3], Nigerian native chicken [4] and Mazandaran indigenous chicken [5]. Great genetic resources embedded in the indigenous poultry await full exploitation that will provide the basis for genetic improvement and diversification to produce breeds that are adapted to local conditions for the benefits of farmers especially in developing countries [4]. The use of major genes to improve productivity in smallholder poultry breeding programs has been researched in various tropical countries (including Indonesia, Malaysia, Thailand, Bangladesh, Bolivia, India, Cameroon, and Nigeria) [6].

The objective of this study was to use the molecular genetics method to validate phenotypic traits-based selection in the *Kambro* chicken line. Phenotypic traits-based is a rapid, practical and cost-saving method which can be applied by breeder especially in the poultry industry as a way to increase the effectiveness of the breeding program. Comb color has a positive significant correlation to sperm function, on the other hand, comb size has a negative significant correlation [7]. These findings were contradictive with other findings which stated that comb size has a positive significant correlation to vitality, sperm function and mating signal in males [8, 9, 10]. Massive amplification of a duplicated sequence in the first intron of *SOX5* as causing the *Pea-comb* phenotype [11]. *Pea-comb* is a dominant mutation in chickens that drastically reduces the size of the comb and wattles [11]. It is an adaptive trait in cold climates as it reduces heat loss and makes the chicken less susceptible to frost lesions [11]. Here we report that *Pea-comb* is caused by a massive amplification of a duplicated sequence located near evolutionary conserved non-coding sequences in intron 1 of the gene encoding the *SOX5* transcription factor [11].

Based on the previous study we hypothesized that by confirming the relation between *SOX5* gene and comb shape can translate into an early screening phase to determine the best male. In this study, the *SOX5* gene polymorphism progenies (BC-III *Kambro*) derived from crossbreed between females *Pelung* and males second backcross generation *Kampong* Broiler-Type (BC-II *Kambro*) were investigated with PCR-RFLP and endonuclease restriction enzyme *HindIII* in the population of *Pea-comb* chicken and *Single-comb* chicken.

Validating this method can help in particular indigenous chicken breeders in Indonesia to be consistent in characterizing and conserving its gene pool.

## 2. Experimental

This study was conducted at the Pusat Inovasi Agro Teknologi (PIAT) Universitas Gadjah Mada, Berbah, Sleman, Daerah Istimewa Yogyakarta, Indonesia. Berbah is situated between latitude 7°47’45.1”S and longitude 110°27’55.0”E at the elevation of 489 m above sea level. PIAT UGM facilitated this study from 2014 until 2019 with the assistance of residents under the supervision of Gama Ayam Research Team, Laboratory of Genetics and Breeding, Faculty of Biology UGM.

This study was performed under the Animal Welfare Act of Indonesia and all procedures involving the handling of animals were approved by the local office of occupational and technical safety (Ethical Clearance Commission of Laboratorium Penelitian dan Pengujian Terpadu, Universitas Gadjah Mada, Yogyakarta No: 00038/04/LPPT/VI/2018).

### 2.1. Experimental animal, DNA isolation and PCR-RFLP

Broodstock consisted of females *Pelung* and males second backcross generation *Kampong* Broiler-Type. Females *Pelung* were acquired from Cianjur, West Java, Indonesia by purchasing from local breeders specializing in *Pelung* breeding with a consistent record of the breeding program. Females *Pelung* and males second backcross generation *Kampong* Broiler-Type were mated in a ratio of 5:1 respectively.

#### 2.1.1. Management of Experimental Birds

Both broodstock and progenies reared under a semi-intensive rearing system with an *ad-libitum* standard feed diet of AD-II and BR-1. Broodstock of each breeding group was fed with *ad-libitum* AD-II (15% Crude Protein) with the administration of the vaccine and prophylactic medications to ensure optimal health of the chickens. Day-Old-Chicks (DOCs) of each breeding group were reared intensively in insulated bamboo pens. DOCs were fed *adlibitum* BR-1 (22% Crude Protein, 3050 Kcal ME/kg). Four-weeks-old chickens of each breeding group then transferred into the larger shed (8m^2^) under a semi-intensive rearing system with an *adlibitum* BR-1 diet for the extents of eight weeks.

#### 2.1.2. Blood Sampling and DNA Isolation

Progenies (third backcross generation *Kampong* Broiler-Type; BC-III *Kambro*) and parental generation (females *Pelung* and males second backcross generation *Kampong* Broiler-Type; BC-II *Kambro*) went through screening phase and then selected for molecular analysis. Both progeny and parental blood were collected with vacutainer (1.5-3 mL) from the axillary vein. Genomic DNA was isolated from a whole blood sample using High Pure PCR Template Preparation Kit Roche. A blood sample (25 µL) was dissolved with 175 µL Phosphate Buffer Saline (PBS). DNA concentration and purity were quantified with the Spark® Reader spectrophotometer (TECAN). DNA later kept dissolved in TE buffer (pH 8.0) and stored in the freezer at −20 ^*o*^C for further use.

#### 2.1.3. Primer Design

Chicken (*Gallus gallus*) *SOX5* gene was acquired from NCBI GenBank with the Ref. Seq. **418195**. Primers used to amplify the *SOX5* gene are `forward primer` (`F): 5`’-AGGTAGCCATGGTGACAAGC-3’`, reverse primer (R):5`’-GATCTGTGAGGCAGCCAGTT-3’. Designed primers produced by Integrated DNA Technologies (IDT) with third-party PT. Genetika Science Indonesia.

#### 2.1.4. PCR-RFLP and Electrophoresis

*The* PCR kit used is Fast Start Roche. PCR-program in amplifying *SOX5* gene consisted of initial denaturation at 92 ^*°*^C for 5 min, final extension at 72 ^*°*^C for 10 min. Cyclic condition to amplify *SOX5* gene fragments consisted of denaturation at 92 ^*°*^C for 1 min, annealing at 48.6 ^*°*^C for 1 min and extension at 72 ^*°*^C for 2 min. The entire cycle consisted of 35 cycles. 10 µL were mixed with ddH2O 18 µL, 10x buffer R 2 µL, and restriction enzyme endonuclease *HindIII* 1 µL. Incubation variants consisted of 2 h, 4 h, 8 h, 12 h, 16 h and 24 h at 37 ^*°*^C.

#### 2.1.5. Electrophoresis

The PCR products and digestion products were detected by electrophoresis with Submarine Electrophoresis System (Mupid-EXU) device. Gel agarose and product ratio were 0.8% isolated DNA, 1.5% PCR products and 2% digestion products in 1x buffer TBE at 50 volts for 30 min. Gel staining with ethidium bromide during absorbance period of 30 min. Finally, observations were performed under ultraviolet light (λ = 260 nm) AnalytikJena^™^ gel imaging system and documented with GelDoc^™^ Documentation System. Images of electrophoresis gel were analyzed with ImageLab (V. 6.0.1) to identify each band based on base-pair length with BenchTop 100 bp ladder.

## 3. Results and Discussion

Based on this study backcross progenies of females *Pelung* and males second backcross generation *Kampong* Broiler-Type (BC-II *Kambro*) were called third backcross generation *Kampong* Broiler-Type (BC-III *Kambro*). The breeding diagram of the development of the third backcross generation *Kambro* chicken line selective breeding program is depicted in Figure 1. Research on *Pelung* chicken genetic pool and phenotypic traits can assist the objective of the *Kampong* Broiler-Type selective breeding program to be the potential meat-producer chicken of Indonesia.

**Figure 1.**
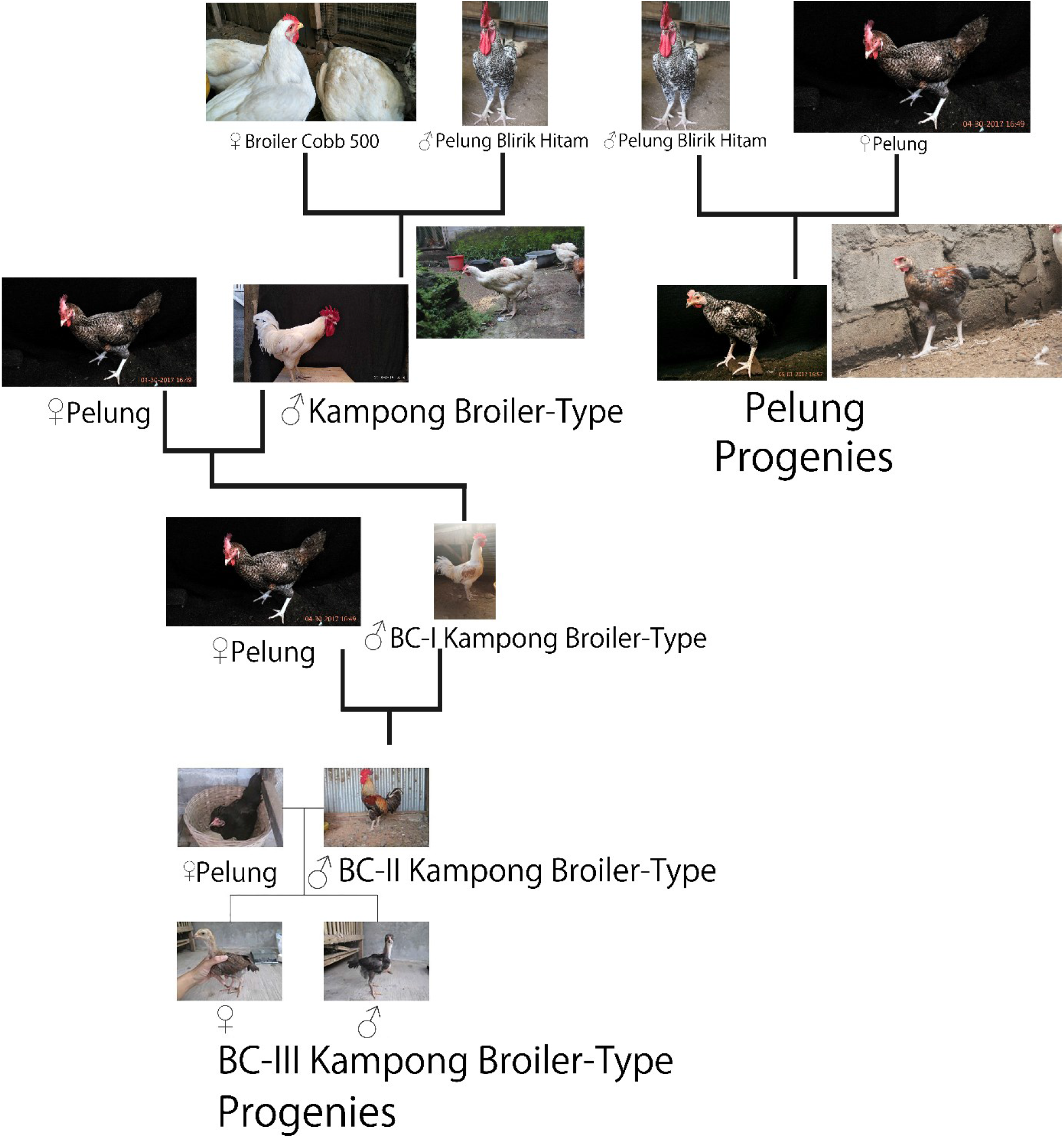
Diagram of the development scheme for third backcross generation *Kambro* chicken line selective breeding program. Crosses between female *Pelung* (P) with male second backcross generation *Kambro* (BC-II *Kambro*) were derived from previous generation of male BC-I *Kambro* and female *Pelung*. BC-I *Kambro* chickens were derived from several cross of main progenies *Kambro* and *Pelung. Kampong* Broiler-Type (*Kambro*) chickens were derived from crossbreed between *Pelung Blirik Hitam* and Broiler Cobb 500 (Mahardhika and Daryono, 2019).

**Figure 2.**
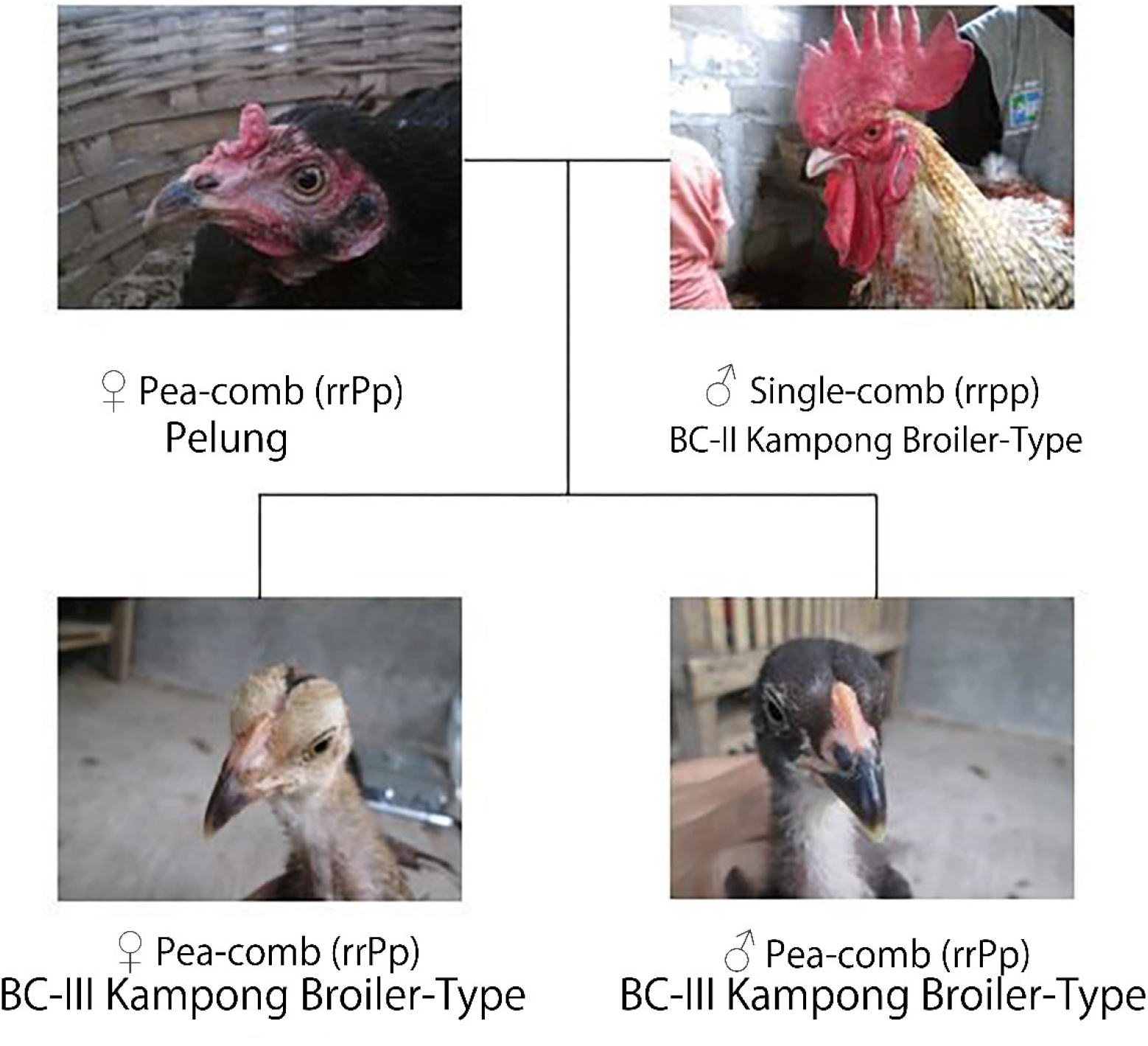
Comb shape of third backcross generation *Kampong* Broiler-Type (*Kambro*) and parentals.

*Kampong* Broiler-Type (Kambro) selective breeding program highlighted five phenotype traits including body weight, shank color, beak color, comb shape and comb color (Table 1). The observation on third backcross generation *Kampong* Broiler-Type (BC-III *Kambro*) indicated allelic inheritance of comb shape and comb color of the parental generation. *Pea-comb* shape is controlled by *rrP_* allele genotype and *Single-comb* shape is controlled by *rrpp* allele genotype. Based on the Mendelian genetics concept, progenies expressing 100% *Pea-comb* shape resulted from the gametic interaction between *rP* and *rp*. Progenies were expressing 50% *Pea-comb* shape and *Single-comb* shape resulted from the gametic interaction between *rP* and *rp* from *rrPp* and *rrpp*.

**Table 1.** Phenotype traits of third backcross generation *Kampong* Broiler-Type (*Kambro*) chickens

| Phenotype Traits | Female *Pelung* | Male BC-II *Kambro* | Female Progenies BC-III *Kambro* | Male Progenies BC-III *Kambro* |
| --- | --- | --- | --- | --- |
| (Comb shape) | (Pea) | (Single) | (Pea) | (Pea) |
| Color | Red | Red | Pale Red | Pale Red |
| Beak | Blackish White | White | Dark Brown | Black |
| Shank | Black | White | White | Black-White |

Female *Pelung* used in this study was derived from an intercross between *Ayam Bangkok* which has *Pea-comb* shape. Following phenotype observation, blood samples were taken to investigate *SOX5* gene polymorphism in *Pea-comb* shape and *Single-comb* shape chickens. DNA isolation from whole blood sample conducted in the Laboratory of Genetics and Breeding, Faculty of Biology, Universitas Gadjah Mada, Yogyakarta, Indonesia. Isolated DNA was quantified with the spectrophotometry method as detailed in Table 2.

**Table 2.** Spectrophotometry result of isolated DNA.

| No. | ID | Absorbancy | Ratio | Concentration (ng/μL) |
| --- | --- | --- | --- | --- |
| 1 | *Anak 1<sub>a</sub>* | 0,011 | 1,959 | 19,8 |
| 2 | *Anak 1<sub>b</sub>* | 0,014 | 2,122 | 22,4 |
| 3 | *Anak 2<sub>a</sub>* | 0,016 | 1,480 | 19,7 |
| 4 | *Anak 2<sub>b</sub>* | 0,018 | 2,064 | 28,3 |
| 5 | Backcross II<sub>a</sub> | 0,018 | 1,387 | 45,5 |
| 6 | Backcross II<sub>b</sub> | 0,041 | 1,951 | 52,9 |
| 7 | *Kampung<sub>a</sub>* | 0,014 | 1,822 | 40,1 |
| 8 | *Kampung<sub>b</sub>* | 0,023 | 1,863 | 47,2 |

Based on spectrophotometry results in Table 2, samples showed concentration ranging from 25 ng/µL to 50 ng/µL. These samples were sufficient to be used as a DNA template for the PCR procedure. PCR products were 503 bp (NCBI GenBank Ref. Seq. **418195**) with annealing temperature was determined through an optimization procedure. The optimization result concluded that 48.6 ^°^C annealing temperature as shown in Figure 3.

**Figure 3.**
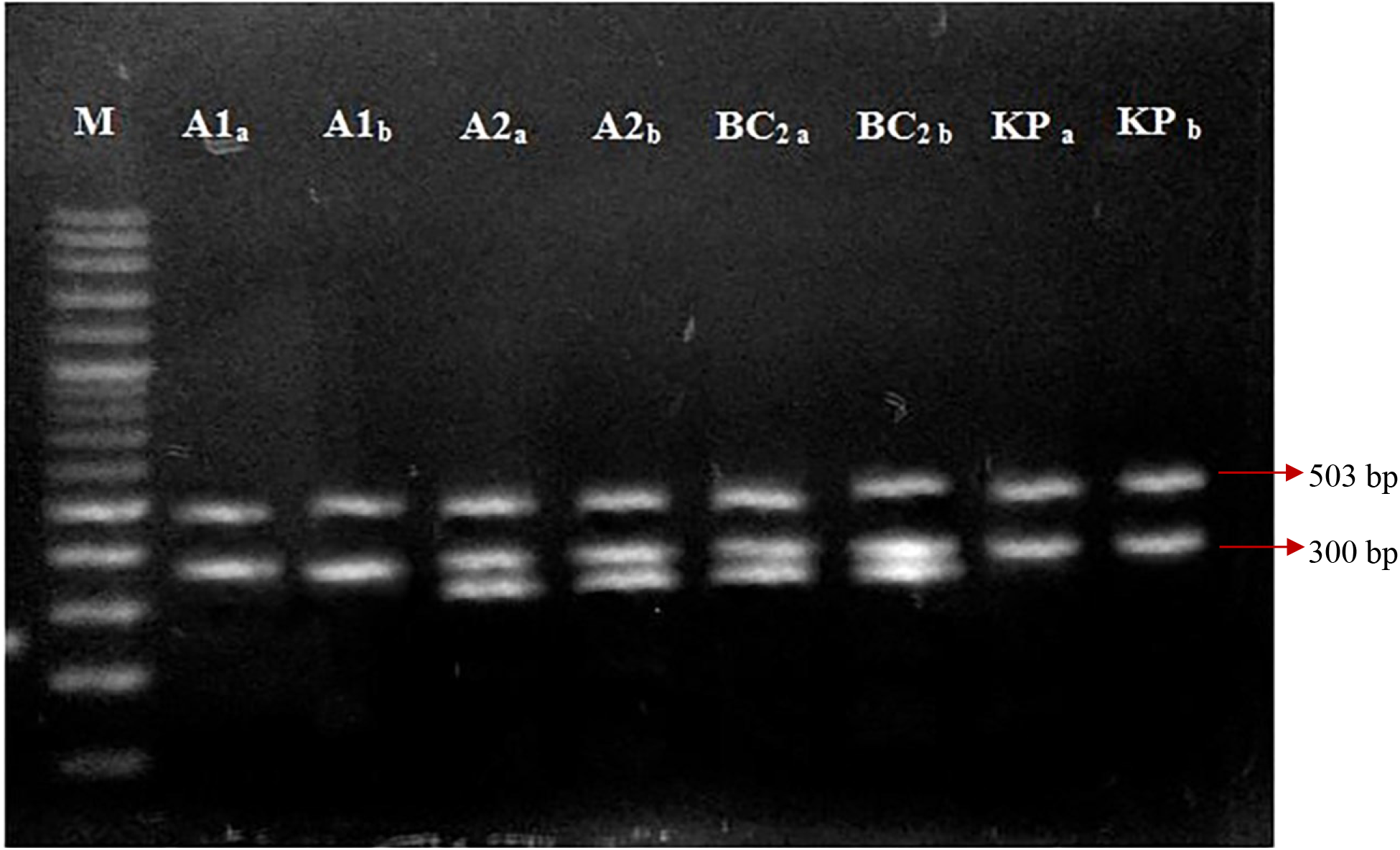
*SOX5* gene PCR visualization. A_1a_: *Anak Ayam 1a*; A_1b_: *Anak Ayam 1b*; A_2a_: *Anak Ayam 2a*; A_2b_: *Anak Ayam 2b*; BC_2a_: backcross II a: BC_2b_: backcross II b; KP_a_: *Ayam kampong a*; KP_b_: *Ayam kampong b*; M: marker DNA 100bp (vivantis)

Figure 3 described all samples to show the target PCR product with a size of 503 bp. Until this step, the *SOX5* gene polymorphism was still indefinable in both *Pea-comb* shape chickens and *Single-comb* shape chickens. Non-target/artifact PCR products with the size of 300 bp were also produced and it required a further optimization step to eliminate this particular product. Non-target/artifact PCR products observed to always appear in temperature ranging from 42 ^°^C to 56 ^°^C. Based on the optimization process it was concluded that the primer used in this study was not specific and recognized another DNA segment in the chicken genome resulted in 300 bp non-target/artifact products.

Following this result endonuclease restriction enzyme digestion performed using *HindIII. HindIII* is a universal restriction enzyme besides *EcoRI* and *BamHI* with restricting point presents in all organisms. Preliminary research was conducted to determine the effectiveness of *HindIII* in recognizing the polymorphism site in the *SOX5* gene sequence. Based on the information required from virtual digest using restriction mapper software GenBank NCBI, *SOX5* gene sequence *Gallus gallus* can be digested with *HindIII* enzyme with the fragmented result of 336 bp and 167 bp.

Figure 4 described the result of chicken (*Gallus gallus*) *SOX5* gene digestion using the *HindIII* endonuclease restriction enzyme. Based on the result the restriction point was not able to produce the expected products. This can be affected by the absence of the restriction point in the DNA sample and contradicts the result of virtual digest using restriction mapper software. This difference can be caused by the difference in *SOX5* gene sequence nucleotide size acquired from GenBank NCBI. Based on this result a sequencing was suggested to confirm the nucleotide structure of DNA samples used.

**Figure 4.**
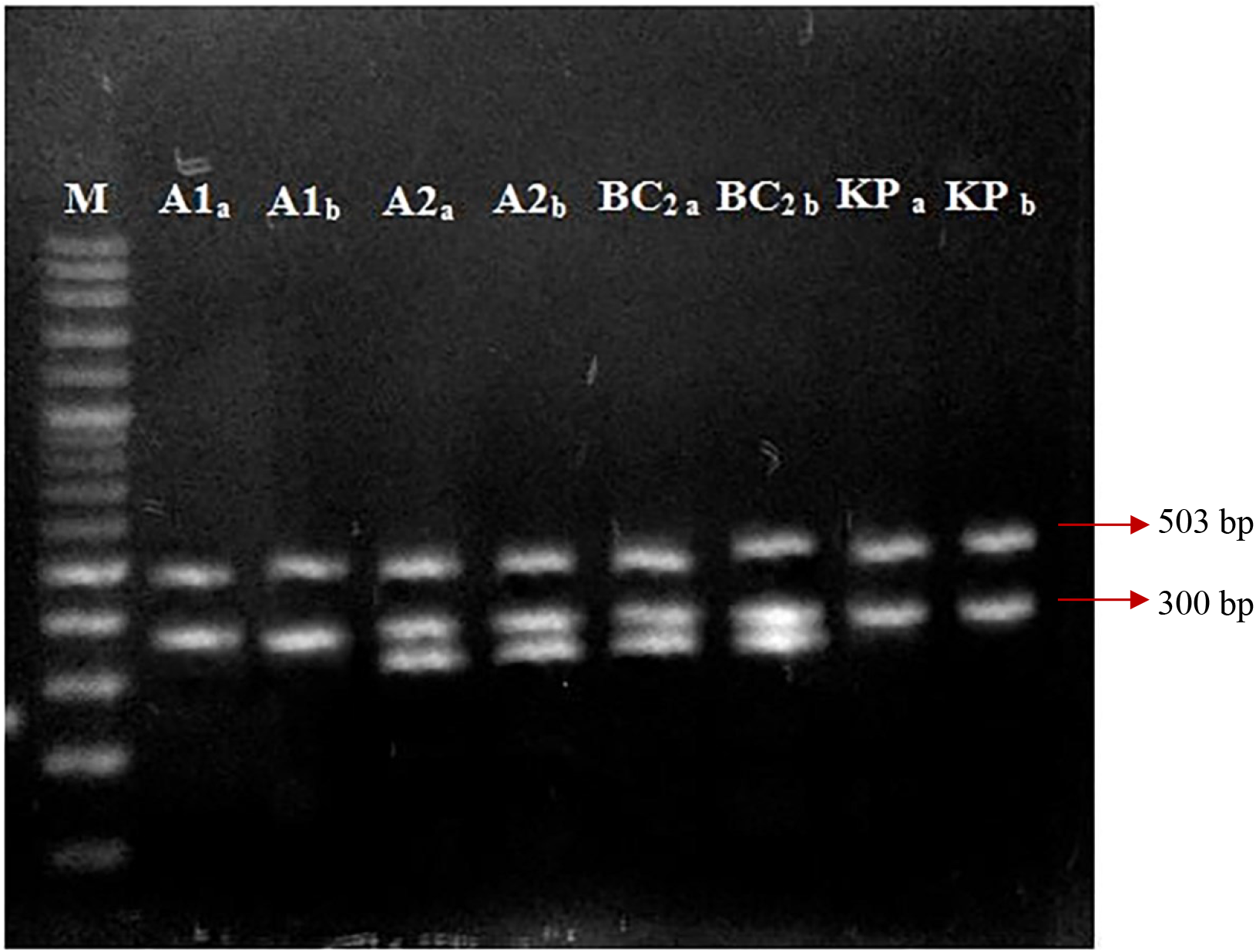
Digestion of chicken (*Gallus gallus*) *SOX5* gene using endonuclease restriction enzyme *HindIII*. A_1a_: *Anak Ayam 1a*; A_1b_: *Anak Ayam 1b*; A_2a_: *Anak Ayam 2a*; A_2b_: *Anak Ayam 2b*; BC_2a_: backcross II a: BC_2b_: backcross II b; KP_a_: *Ayam kampong a*; KP_b_: *Ayam kampong b*; M: marker DNA 100bp (vivantis)

Another factor affected the results of this study was sample preservation and incubation time. Incubation period variations performed in this study were classified into 5 varieties, 2 h, 4 h, 8 h, 16 h, and 24 h. Based on the result different incubation periods showed a negative correlation on the digestion results. The effective incubation period during the digestion step using restriction enzyme *HindIII* was around 4 h to 5 h [12]. *SOX5* gene in this study was unable to be digested using endonuclease restriction enzyme *HindIII*.

## 4. Conclusion

The result of *SOX5* gene polymorphism showed the comb shape uniformity between progenies and parental of *Kambro* backcross generation. This study concluded that the genotype of *Pea-comb* shape female *Pelung* was undetermined and there was no polymorphism of the *SOX5* gene between pea and single comb. PCR-RFLP using endonuclease restriction enzyme *HindIII* produced both target products and non-target/artifact products. The sequencing procedure was required to provide nucleotide sequences. The molecular genetics method to validate phenotypic traits-based selection in the *Kambro* chicken line can be implemented as an early screening phase to select the best male. Further improvement and alternative ways can be applied including sequencing-based selection.

## Supporting information

Experimental File

## References

[1] Mahardhika IWS & Daryono BS 2019 Bul. Vet. 11(2) 188–202

[2] Henuk YL & Bakti D 2018 Benefits of Promoting Native Chickens for Sustainable Rural Poultry Development in Indonesia Mohammad Basyuni, S. Hut. M.Si., Ph.D., Prof. Dr. Ir. Elisa Julianti, M.Si [editors] Conference Proceeding of Seminar Ilmiah Nasional Dies Natalis USU-64 Sumatera Utara [Indonesia] University of Sumatera Utara 69–76

[3] Manjula P, Park H-B, Seo D, Choi N, Jin S, Ahn SJ, Heo KN, Kang BS and Lee J-H 2018 Asian-Australas J. Anim. Sci. 31(1) 26–31

[4] Nwenya JMI, Nwakpu EP, Nwose RN and Ogbuagu KP 2017 J. Poult. Res. 14(2) 07–11

[5] Niknafs Sh, Abdi H, Fatemi SA, Zandi MB and Baneh H 2013 J. Bio. 3(1) 25–31

[6] FAO 2004 Small-scale poultry production: technical guide (FAO Animal Production and Health)

[7] Navara KJ, Anderson EM and Edwards ML 2012 Behav. Eco. Adv. Acc. 12 1036–1041

[8] Gebriel GM, Kalamah M, El-Fiky AA and Ali AFA 2009 Egypt Poult. Sci. 29 677–693

[9] El Ghany FAA, El Dein A, Soliman MM, Rezaa AM and El Sodany S M 2011 Egypt Poult. Sci. 31 331–349

[10] Udeh I, Ugwu SOC and Ogagifo NL 2011 Asian J. Anim. Sci. 5 268–276

[11] Wright D, Boije H, Meadows JRS, Bed’hom B, Gourichon D, Vieaud A, Tixier-Boichard M, Rubin C-J, Imsland F, Hallbook F and Andersson L 2009 PLoS Gen. 5(6) e1000512

[12] Dubey A K, Hussain N and Mittal N 2010 J. Nat. Sci. Biol. Med. 1(1) 25–28

