## Supplementary material for "Third Backcross Generation Indonesian Indigenous Chicken *Kampong* Broiler-Type (*Kambro*) *SOX5* Gene Polymorphism": Experimental File

### Experimental File by Mahardhika, IWS

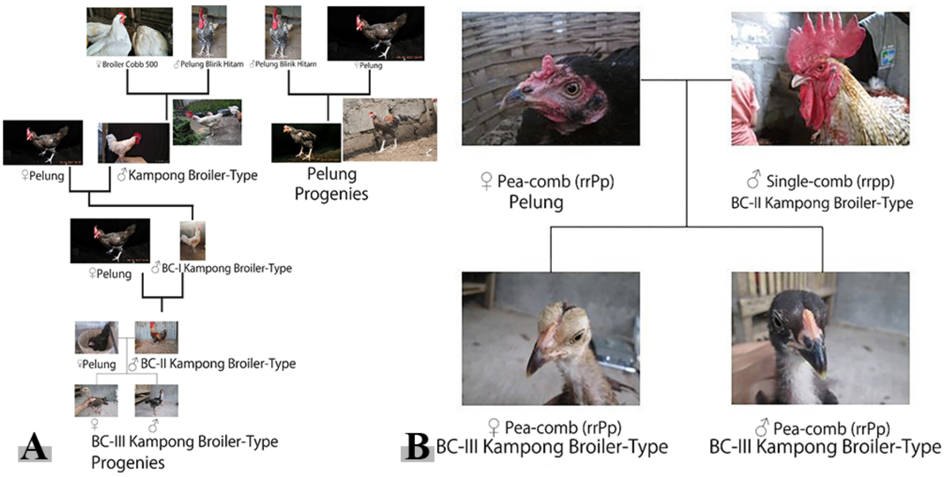


**Fig. 1.** Selective breeding scheme for Indonesia's exotic indigenous chicken and *Kampong* Broiler-type chicken. **A**) Crossbreeding of *Pelung* hens and second backcross generation *Kambro* (BC-II *Kambro*) roosters was the latest breeding program. BC-II *Kambro* roosters were progenies of the previous crossbreeds of *Pelung* hens and first backcross generation *Kambro* (BC-I *Kambro*) roosters. BC-I *Kambro* roosters were progenies of the previous crossbreeds of *Pelung* hens and *Kampong* Broiler-type (*Kambro*) roosters (Perdamaian et al. 2017). *Kambro* roosters were progenies of the previous crossbreeds between *Pelung Blirik Hitam* and Broiler Cobb 500 (Mahardhika and Daryono 2019). **B**) The representative scheme of *Pea-comb Pelung* hen and *Single-comb* BC-II *Kambro* rooster produced third backcross generation *Kambro* (BC-III *Kambro*) progenies with *Pea-comb* shape and *Single-comb* shape (not shown) in both hen and rooster.

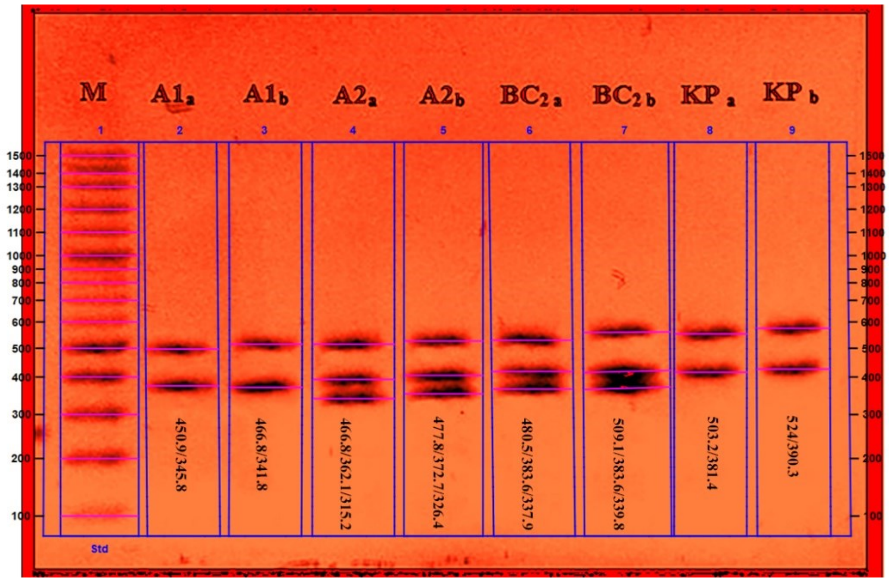


**Fig. 2.** Digestion products of the *Gallus gallus SOX 5* gene from eight representative chickens. Amplified DNA was digested with *HindIII* and separated by electrophoresis on a 2% agarose gel. Lane M is the molecular mass standard of VC 100 bp Plus DNA ladder (Vivantis Technologies). Lane 2 until 9 represents each sample as the following $A_{1a-2a}$: BC-III *Kambro* hen, $A_{1b-2b}$: BC-III *Kambro* rooster, $KP_{a-b}$: *Pelung* hen, $BC_{2a-sb}$: BC-II *Kambro* rooster. Fragment length measurement based on the gel image analysis using ImageLab (V. 6.0.1) under Ethidium Bromide (EtBr) image colors mode adjustment. Molecular weight analysis standard used ImageLab 6.0.1 Bio-Rad 100 bp PCR Molecular Ruler with linear (semi-log) regression method.

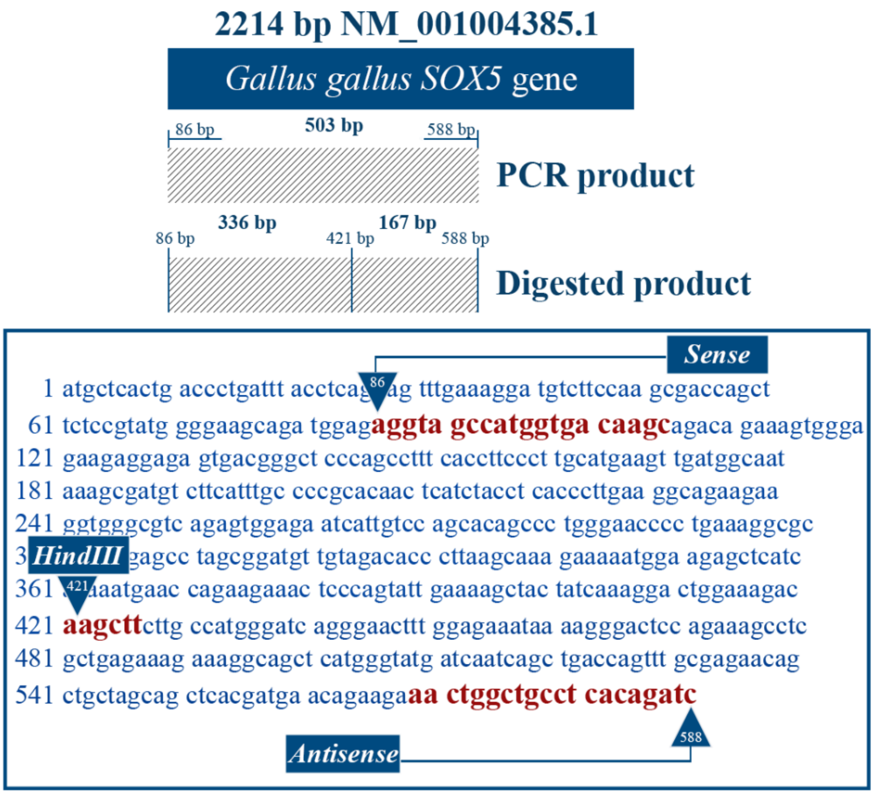


**Fig. 3.** The primers and restriction site of 2214 bp *Gallus gallus SOX 5* gene. The PCR-amplified region corresponding to nucleotide position 86 to 588 of 2214 bp **NM_001004385.1** accession **XM_416425** is indicated as the hatched bar. Primers start from the sense region and antisense region respectively at positions 86 and 588 produced the expected amplification product length of 503 bp. The digested region corresponding to the amplified region consists of two products with different lengths, 336 bp and 167 bp with the *HindIII* palindromic restriction site 5' AAGCTT 3' at nucleotide position 421 of 2214 bp **NM_001004385.1** accession **XM_416425**.

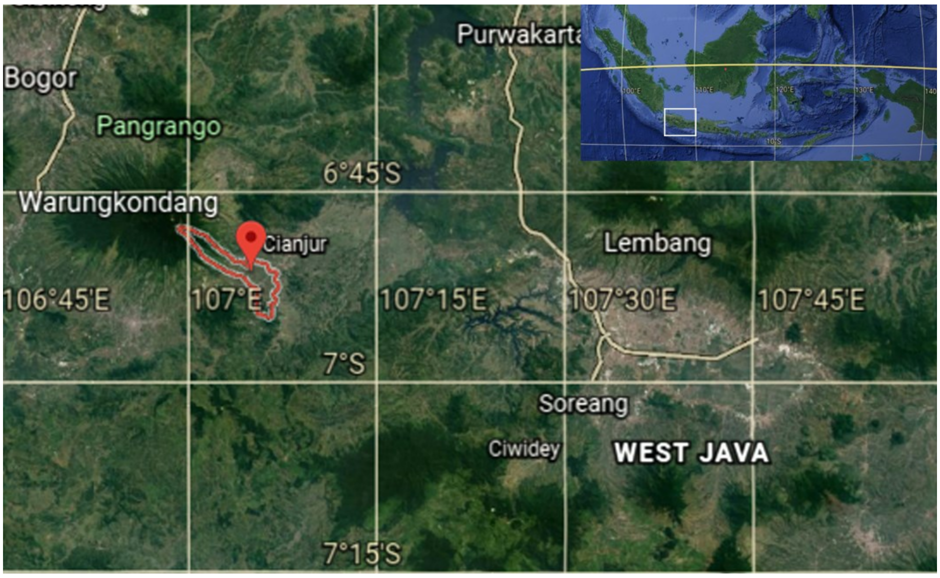


**Fig. 4.** The satellite image of Warungkondang, Cianjur, West Java the region in which *Pelung* chicken originated. *Pelung* or locally named *Ayam Kampong* was first documented and identified from the region which is known as Warungkondang village in 1850 and continues up until the present days. The image was taken from Google Earth Version 9.125.0.0- https://earth.google.com/web

**Table 1.** Typing by PCR-RFLP with *HindIII* and comparison of the fragments with *Gallus gallus SOX5* gene reference sequence (**NM_001004385.1** accession **XM_416425**) (see also Figure 2)

| Chicken ID (lanes, bands) | Fragment lengths (bp) | Product length (bp) | Calculated from | Restriction site location | Nucleotide position |
|---|---|---|---|---|---|
| Hens | | | | | |
| $A_{1a}$ (2, 2) | 451/346 | 346 | Gel | 1 | 536/537 |
| $A_{2a}$ (4, 2) | 467/362/315 | 677 | Gel | 2 | 552/553, 915/916 |
| $BC_{2a}$ (6, 3) | 481/384/338 | 722 | Gel | 2 | 556/557, 951/952 |
| $KP_{a}$ (8, 3) | 503/381 | 381 | Gel | 1 | 466/467 |
| Roosters | | | | | |
| $A_{1b}$ (3, 2) | 467/342 | 342 | Gel | 1 | 552/553 |
| $A_{2b}$ (5, 3) | 478/373/326 | 699 | Gel | 2 | 563/564, 937/938 |
| $BC_{2b}$ (7, 3) | 509/384/340 | 724 | Gel | 2 | 594/595, 979/980 |
| $KP_{b}$ (9, 2) | 524/390 | 390 | Gel | 1 | 609/610 |
| Ref Seq **NM_001004385.1** | 336/167 | 503 | Nuc Seq | 1 | 421/422 |

$A_{1a-2a}$: BC-III *Kambro* hen, $A_{1b-2b}$: BC-III *Kambro* rooster, $KP_{a-b}$: *Pelung* hen, $BC_{2a-sb}$: BC-II *Kambro* rooster. Fragment length measurement based on the gel image analysis using ImageLab (V. 6.0.1) under Ethidium Bromide (EtBr) image colors mode adjustment. Molecular weight analysis standard used ImageLab 6.0.1 Bio-Rad 100 bp PCR Molecular Ruler with linear (semi-log) regression method.

**Table 2.** Quantification results from isolated and purified chicken DNA extracted from the whole blood sample

| Chicken ID (n= 8) | Absorbency | Ratio ($\lambda_{260/280}$) | Concentration (ng/µL) |
|---|---|---|---|
| $A_{1a}$ | 0.011 | 1.959 | 19.8 |
| $A_{1b}$ | 0.014 | 2.122 | 22.4 |
| $A_{2a}$ | 0.016 | 1.480 | 19.7 |
| $A_{2b}$ | 0.018 | 2.064 | 28.3 |
| $BC_{2a}$ | 0.018 | 1.387 | 45.5 |
| $BC_{2b}$ | 0.041 | 1.951 | 52.9 |
| $KP_{a}$ | 0.014 | 1.822 | 40.1 |
| $KP_{b}$ | 0.023 | 1.863 | 47.2 |

$A_{1a-2a}$: BC-III *Kambro* hen, $A_{1b-2b}$: BC-III *Kambro* rooster, $KP_{a-b}$: *Pelung* hen, $BC_{2a-sb}$: BC-II *Kambro* rooster. Spectrophotometry used $\lambda_{260/280}$ Spark® Reader UV-Vis spectrophotometer (TECAN).
