## Supplemental File for "Third Backcross Generation Indonesian Indigenous Chicken *Kampong* Broiler-Type (*Kambro*) *SOX5* Gene Polymorphism"

### Image Report: RFLP

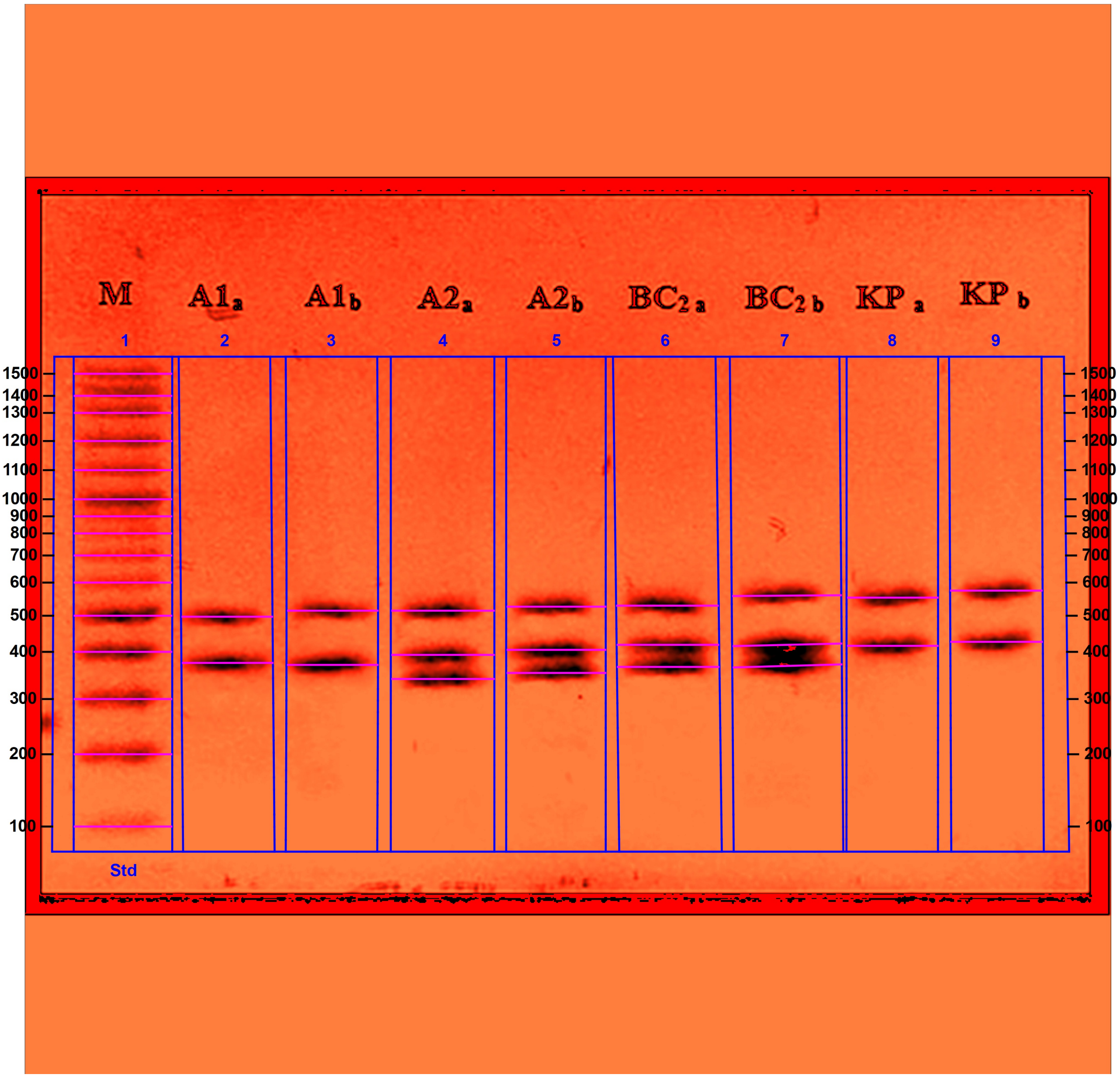


D:\PASCASARJANA\Tesis\JURNAL_2020\ABS3389\RFLP.scn

#### Acquisition Information

| | |
|---|---|
| Program | Adobe Photoshop CC 2019 (Windows) |

#### Image Information

| | |
|---|---|
| Acquisition Date | 12/22/2020 6:54:46 PM |
| User Name | I Wayan Swarautama M |
| Image Area (mm) | X: 91.4 Y: 91.4 |
| Pixel Size (µm) | X: 84.7 Y: 84.7 |

| Data Range (Int) | 0 - 255 |
|---|---|

#### Analysis Settings

| | |
|---|---|
| Detection | Lane detection:<br>Automatically detected lanes with manual adjustments<br><br>Band detection:<br>Automatically detected bands with sensitivity: High<br>Manually adjusted bands<br><br>Lane Background Subtraction:<br>Lane background subtracted with disk size: 0.1<br><br>Lane width: Variable |
| Mol. Weight Analysis | Standard: Bio-Rad 100 bp PCR Molecular Ruler<br>Standard lanes: first<br>Regression method: Linear (semi-log) |

#### Lane Statistics

| Lane No. | Adj. Total Band Vol. (Int) | Total Band Vol. (Int) | Adj. Total Lane Vol. (Int) | Total Lane Vol. (Int) | Bkgd. Vol. (Int) | Norm. Factor |
|---|---|---|---|---|---|---|
| 1 | 677,572 | 1,715,784 | 681,786 | 1,882,580 | 1,200,794 | N/A |
| 2 | 263,809 | 360,906 | 268,359 | 985,257 | 716,898 | N/A |
| 3 | 294,021 | 367,185 | 297,752 | 882,882 | 585,130 | N/A |
| 4 | 322,802 | 520,974 | 325,480 | 1,102,100 | 776,620 | N/A |
| 5 | 332,937 | 541,332 | 338,877 | 1,070,388 | 731,511 | N/A |
| 6 | 318,138 | 557,736 | 321,300 | 1,088,544 | 767,244 | N/A |
| 7 | 300,404 | 623,262 | 304,110 | 1,105,587 | 801,477 | N/A |
| 8 | 273,000 | 337,064 | 275,639 | 612,066 | 336,427 | N/A |
| 9 | 249,228 | 283,268 | 250,792 | 461,196 | 210,404 | N/A |

#### Lane And Band Analysis

##### Lane 1 - Bio-Rad 100 bp PCR Molecular Ruler

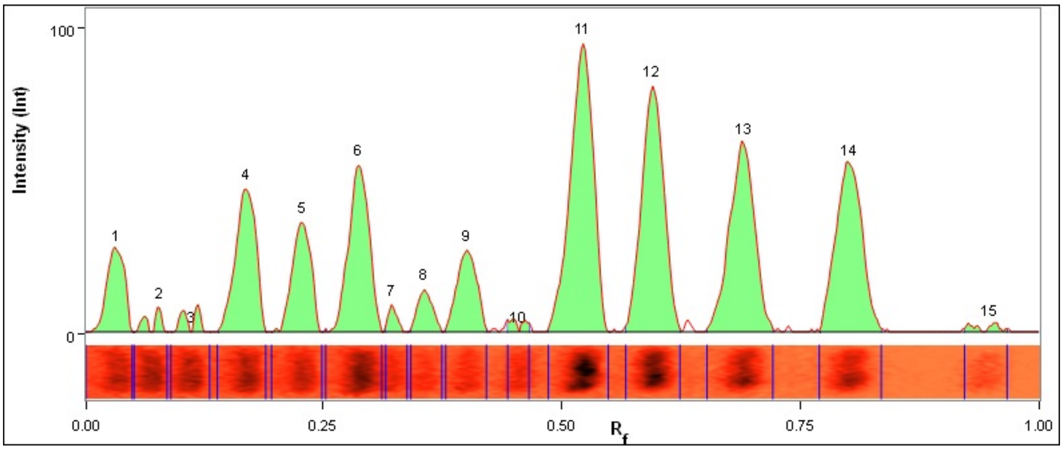


| Lane | Band No. | Band Label | Base Pairs (bp) | Relative Front | Adj. Volume (Int) | Volume (Int) | Abs. Quant. | Rel. Quant. | Band % | Lane % |
|---|---|---|---|---|---|---|---|---|---|---|
| 1 | 1 | | 1,500.0 | 0.034 | 29,302 | 129,360 | N/A | N/A | 4.3 | 4.3 |
| 1 | 2 | | 1,400.0 | 0.079 | 4,998 | 112,112 | N/A | N/A | 0.7 | 0.7 |
| 1 | 3 | | 1,300.0 | 0.114 | 6,370 | 118,874 | N/A | N/A | 0.9 | 0.9 |

| | | | | | | | | | | |
|---|---|---|---|---|---|---|---|---|---|---|
| 1 | 4 | | 1,200.0 | 0.170 | 57,526 | 141,610 | N/A | N/A | 8.5 | 8.4 |
| 1 | 5 | | 1,100.0 | 0.229 | 40,376 | 127,792 | N/A | N/A | 6.0 | 5.9 |
| 1 | 6 | | 1,000.0 | 0.288 | 66,444 | 184,632 | N/A | N/A | 9.8 | 9.7 |
| 1 | 7 | | 900.0 | 0.323 | 4,606 | 53,998 | N/A | N/A | 0.7 | 0.7 |
| 1 | 8 | | 800.0 | 0.357 | 12,446 | 63,700 | N/A | N/A | 1.8 | 1.8 |
| 1 | 9 | | 700.0 | 0.402 | 31,948 | 81,536 | N/A | N/A | 4.7 | 4.7 |
| 1 | 10 | | 600.0 | 0.456 | 3,430 | 52,332 | N/A | N/A | 0.5 | 0.5 |
| 1 | 11 | | 500.0 | 0.523 | 129,164 | 191,394 | N/A | N/A | 19.1 | 18.9 |
| 1 | 12 | | 400.0 | 0.596 | 106,624 | 161,308 | N/A | N/A | 15.7 | 15.6 |
| 1 | 13 | | 300.0 | 0.692 | 92,904 | 142,198 | N/A | N/A | 13.7 | 13.6 |
| 1 | 14 | | 200.0 | 0.803 | 88,102 | 118,874 | N/A | N/A | 13.0 | 12.9 |
| 1 | 15 | Reference Band | 100.0 | 0.949 | 3,332 | 36,064 | N/A | N/A | 0.5 | 0.5 |

| | |
|---|---|
| Band Detection | Automatically detected bands with sensitivity: High |
| Lane Background | Lane background subtracted with disk size: 0.1 |
| Lane Width | 8.30 mm |
| Regression Equation | y = -1.24 * x + 3.3<br>R-squared value: R^2 = 0.972429 |

#### Lane 2

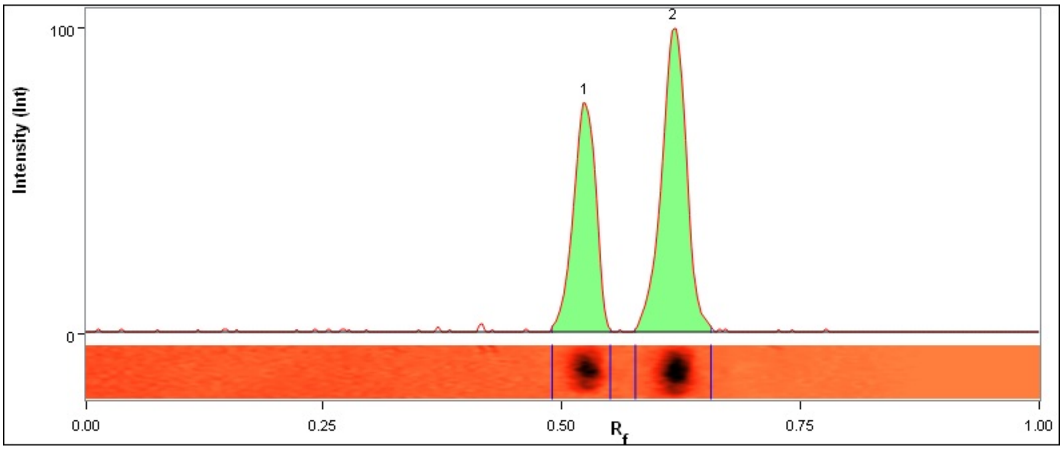


| Lane | Band No. | Band Label | Base Pairs (bp) | Relative Front | Adj. Volume (Int) | Volume (Int) | Abs. Quant. | Rel. Quant. | Band % | Lane % |
|---|---|---|---|---|---|---|---|---|---|---|
| 2 | 1 | | 450.9 | 0.525 | 106,652 | 154,609 | N/A | N/A | 40.4 | 39.7 |
| 2 | 2 | | 345.8 | 0.619 | 157,157 | 206,297 | N/A | N/A | 59.6 | 58.6 |

| | |
|---|---|
| Band Detection | Automatically detected bands with sensitivity: High |
| Lane Background | Lane background subtracted with disk size: 0.1 |
| Lane Width | 7.70 mm |
| Regression Equation | y = -1.24 * x + 3.3<br>R-squared value: R^2 = 0.972429 |

#### Lane 3

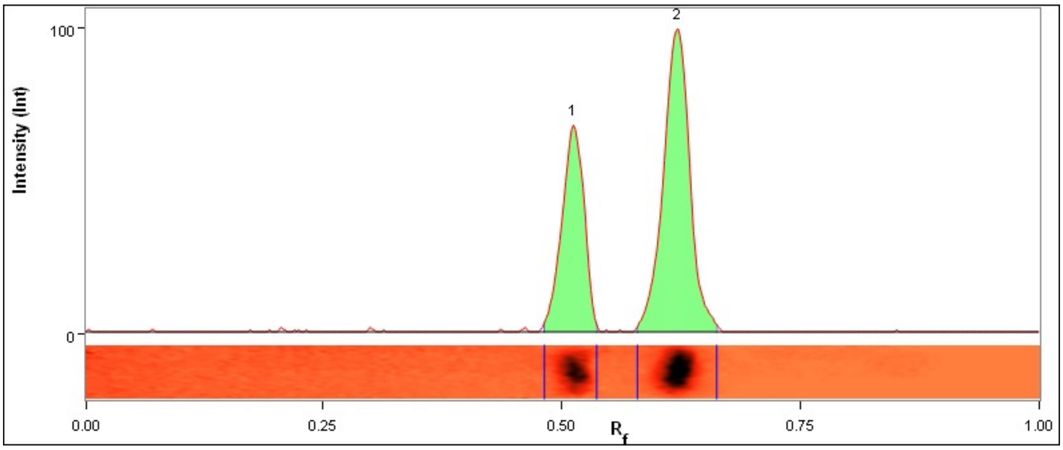


| Lane | Band No. | Band Label | Base Pairs (bp) | Relative Front | Adj. Volume (Int) | Volume (Int) | Abs. Quant. | Rel. Quant. | Band % | Lane % |
|---|---|---|---|---|---|---|---|---|---|---|
| 3 | 1 | | 466.8 | 0.513 | 107,198 | 146,328 | N/A | N/A | 36.5 | 36.0 |
| 3 | 2 | | 341.8 | 0.623 | 186,823 | 220,857 | N/A | N/A | 63.5 | 62.7 |

| | |
|---|---|
| Band Detection | Automatically detected bands with sensitivity: High |
| Lane Background | Lane background subtracted with disk size: 0.1 |
| Lane Width | 7.70 mm |
| Regression Equation | y = -1.24 * x + 3.3<br>R-squared value: R^2 = 0.972429 |

#### Lane 4

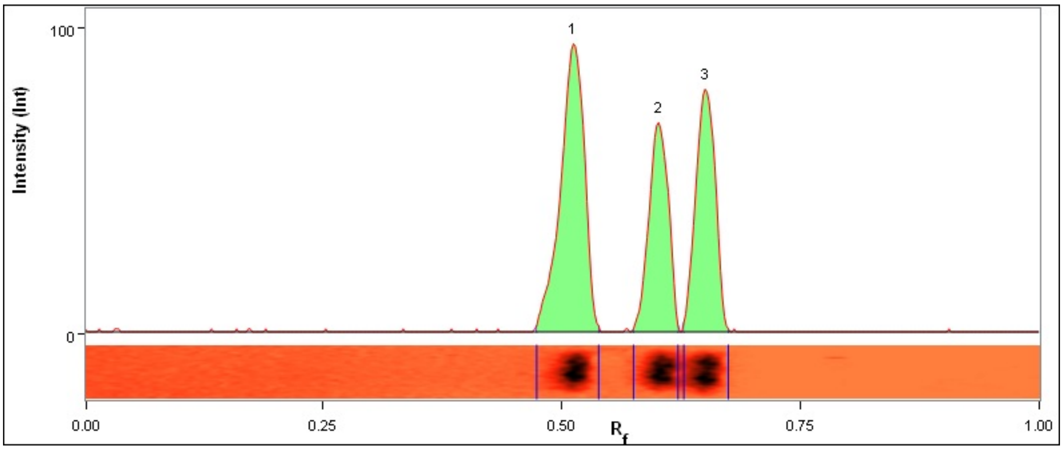


| Lane | Band No. | Band Label | Base Pairs (bp) | Relative Front | Adj. Volume (Int) | Volume (Int) | Abs. Quant. | Rel. Quant. | Band % | Lane % |
|---|---|---|---|---|---|---|---|---|---|---|
| 4 | 1 | | 466.8 | 0.513 | 141,110 | 187,769 | N/A | N/A | 43.7 | 43.4 |
| 4 | 2 | | 362.1 | 0.602 | 83,842 | 170,671 | N/A | N/A | 26.0 | 25.8 |
| 4 | 3 | | 315.2 | 0.651 | 97,850 | 162,534 | N/A | N/A | 30.3 | 30.1 |

| | |
|---|---|
| Band Detection | Automatically detected bands with sensitivity: High |
| Lane Background | Lane background subtracted with disk size: 0.1 |
| Lane Width | 8.72 mm |
| Regression Equation | y = -1.24 * x + 3.3<br>R-squared value: R^2 = 0.972429 |

#### Lane 5

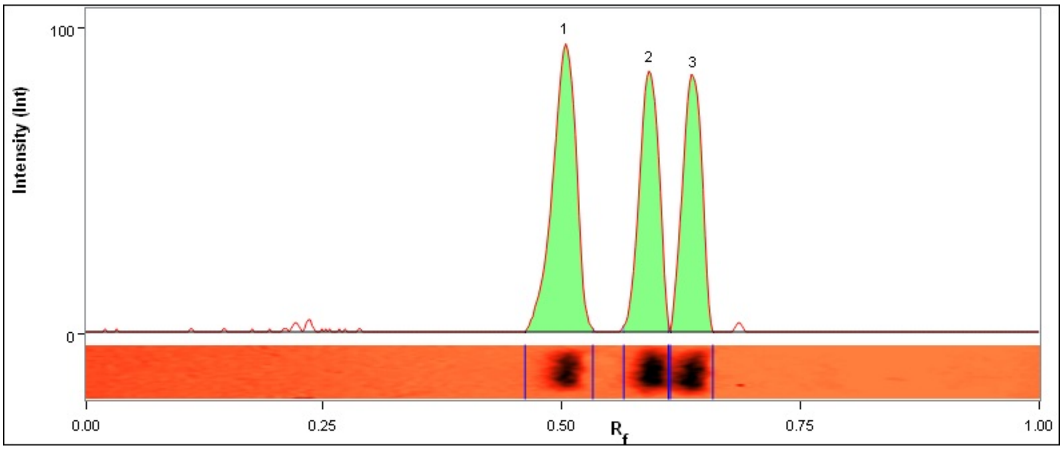


| Lane | Band No. | Band Label | Base Pairs (bp) | Relative Front | Adj. Volume (Int) | Volume (Int) | Abs. Quant. | Rel. Quant. | Band % | Lane % |
|---|---|---|---|---|---|---|---|---|---|---|
| 5 | 1 | | 477.8 | 0.505 | 133,749 | 169,191 | N/A | N/A | 40.2 | 39.5 |
| 5 | 2 | | 372.7 | 0.592 | 101,079 | 198,792 | N/A | N/A | 30.4 | 29.8 |
| 5 | 3 | | 326.4 | 0.639 | 98,109 | 173,349 | N/A | N/A | 29.5 | 29.0 |

| | |
|---|---|
| Band Detection | Automatically detected bands with sensitivity: High |
| Lane Background | Lane background subtracted with disk size: 0.1 |
| Lane Width | 8.38 mm |
| Regression Equation | y = -1.24 * x + 3.3<br>R-squared value: R^2 = 0.972429 |

#### Lane 6

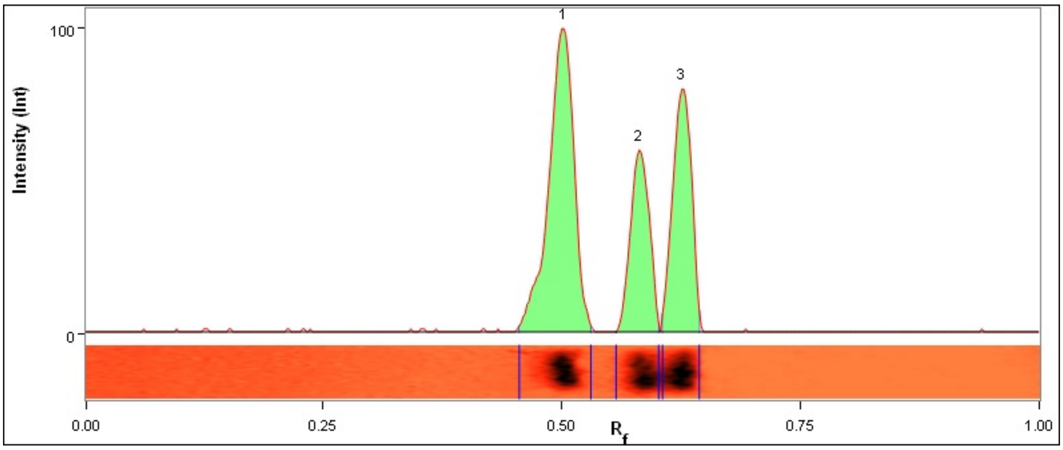


| Lane | Band No. | Band Label | Base Pairs (bp) | Relative Front | Adj. Volume (Int) | Volume (Int) | Abs. Quant. | Rel. Quant. | Band % | Lane % |
|---|---|---|---|---|---|---|---|---|---|---|
| 6 | 1 | | 480.5 | 0.503 | 155,040 | 208,896 | N/A | N/A | 48.7 | 48.3 |
| 6 | 2 | | 383.6 | 0.582 | 69,768 | 179,010 | N/A | N/A | 21.9 | 21.7 |
| 6 | 3 | | 337.9 | 0.627 | 93,330 | 169,830 | N/A | N/A | 29.3 | 29.0 |

| | |
|---|---|
| Band Detection | Automatically detected bands with sensitivity: High |
| Lane Background | Lane background subtracted with disk size: 0.1 |

| | |
|---|---|
| Lane Width | 8.64 mm |
| Regression Equation | y = -1.24 * x + 3.3<br>R-squared value: R^2 = 0.972429 |

##### Lane 7

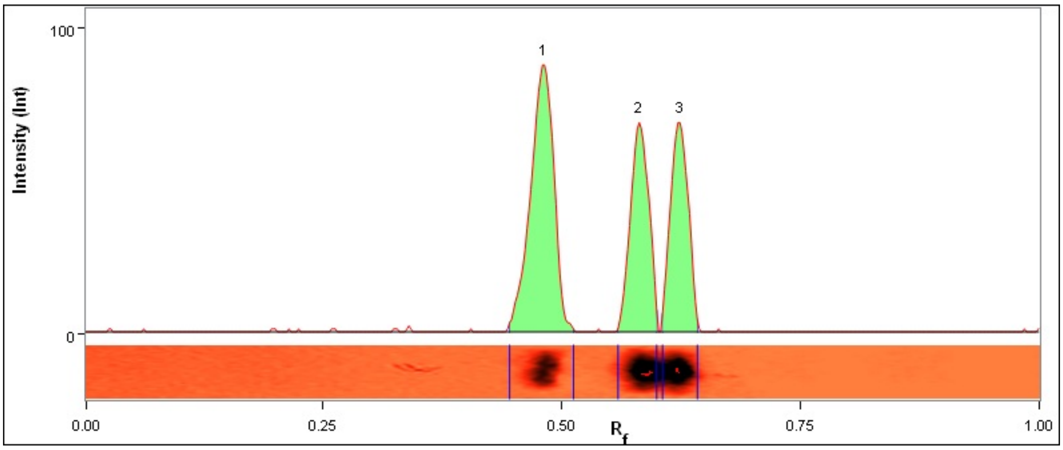


| Lane | Band No. | Band Label | Base Pairs (bp) | Relative Front | Adj. Volume (Int) | Volume (Int) | Abs. Quant. | Rel. Quant. | Band % | Lane % |
|---|---|---|---|---|---|---|---|---|---|---|
| 7 | 1 | | 509.1 | 0.483 | 135,160 | 165,898 | N/A | N/A | 45.0 | 44.4 |
| 7 | 2 | | 383.6 | 0.582 | 83,821 | 247,975 | N/A | N/A | 27.9 | 27.6 |
| 7 | 3 | | 339.8 | 0.625 | 81,423 | 209,389 | N/A | N/A | 27.1 | 26.8 |

| | |
|---|---|
| Band Detection | Automatically detected bands with sensitivity: High |
| Lane Background | Lane background subtracted with disk size: 0.1 |
| Lane Width | 9.23 mm |
| Regression Equation | y = -1.24 * x + 3.3<br>R-squared value: R^2 = 0.972429 |

##### Lane 8

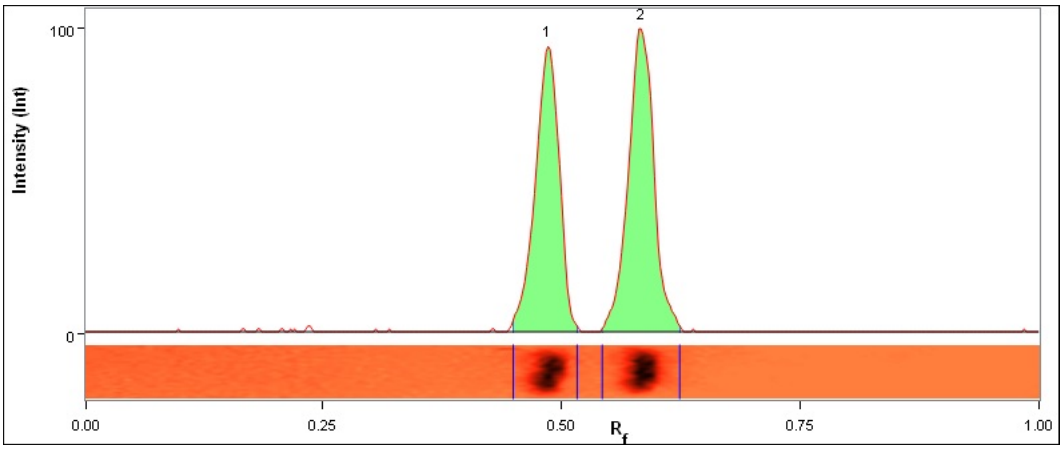


| Lane | Band No. | Band Label | Base Pairs (bp) | Relative Front | Adj. Volume (Int) | Volume (Int) | Abs. Quant. | Rel. Quant. | Band % | Lane % |
|---|---|---|---|---|---|---|---|---|---|---|
| 8 | 1 | | 503.2 | 0.487 | 127,855 | 160,251 | N/A | N/A | 46.8 | 46.4 |
| 8 | 2 | | 381.4 | 0.584 | 145,145 | 176,813 | N/A | N/A | 53.2 | 52.7 |

| | |
|---|---|
| Band Detection | Automatically detected bands with sensitivity: High |
| Lane Background | Lane background subtracted with disk size: 0.1 |
| Lane Width | 7.70 mm |
| Regression Equation | y = -1.24 * x + 3.3<br>R-squared value: R^2 = 0.972429 |

#### Lane 9

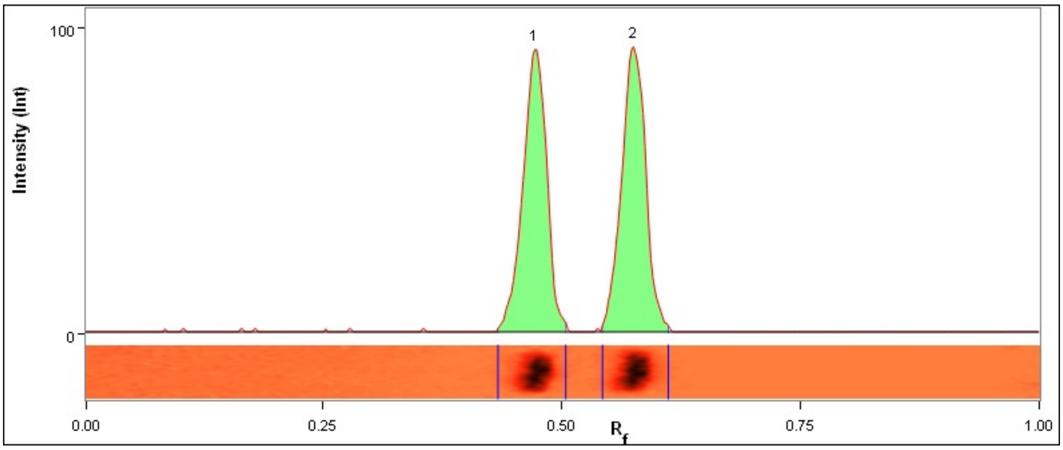


| Lane | Band No. | Band Label | Base Pairs (bp) | Relative Front | Adj. Volume (Int) | Volume (Int) | Abs. Quant. | Rel. Quant. | Band % | Lane % |
|---|---|---|---|---|---|---|---|---|---|---|
| 9 | 1 | | 524.0 | 0.473 | 122,912 | 139,472 | N/A | N/A | 49.3 | 49.0 |
| 9 | 2 | | 390.3 | 0.576 | 126,316 | 143,796 | N/A | N/A | 50.7 | 50.4 |

| | |
|---|---|
| Band Detection | Automatically detected bands with sensitivity: High |
| Lane Background | Lane background subtracted with disk size: 0.1 |
| Lane Width | 7.79 mm |
| Regression Equation | y = -1.24 * x + 3.3<br>R-squared value: R^2 = 0.972429 |
